# Whole genome analysis for 163 guide RNAs in Cas9 edited mice reveals minimal off-target activity

**DOI:** 10.1101/2021.08.11.455876

**Authors:** Kevin A. Peterson, Sam Khalouei, Joshua A. Wood, Denise G. Lanza, Lauri G. Lintott, Brandon J. Willis, John R. Seavitt, Nour Hanafi, Robert E. Braun, Mary E. Dickinson, Jacqueline K. White, K.C. Kent Lloyd, Jason D. Heaney, Stephen A. Murray, Arun Ramani, Lauryl M.J. Nutter

## Abstract

The Knockout Mouse Phenotyping Program (KOMP^2^) uses CRISRPR/Cas9 for high-throughput mouse line production to generate null alleles in the inbred C57BL/6N strain for broad-based *in vivo* phenotyping. In order to assess the risk of spurious *S. pyogenes* Cas9-induced off-target mutagenesis, we applied whole genome sequencing to compare the genomes of 50 Cas9-derived founder mice representing 163 different gRNAs to 28 untreated inbred control mice. Our analysis pipeline detected 28 off-target sequence variants associated with 21 guides. These potential off-targets were identified in 18/50 (36%) founders with 9/28 (32%) independently validated corresponding to 8 founder animals. In total, only 4.9% (8/163) of all guides exhibited off-target activity resulting in a rate of 0.16 Cas9 off-target mutations per founder analyzed. In comparison, we observed ~1225 unique variants in each mouse regardless of whether or not it was exposed to Cas9. These findings indicate that Cas9-mediated off-target mutagenesis is rare in founder knockout mice generated using guide RNAs designed to minimize off-target risk. Overall, bona fide off-target variants comprise a small fraction of the genetic heterogeneity found in carefully maintained colonies of inbred strains.

## Main

CRISPR Cas9 genome editing has tremendous therapeutic potential for treating a large number of human diseases^1^. The widely used *S. pyogenes* Cas9 is a programmable RNA-guided endonuclease that can be targeted to precise locations in the genome of virtually any organism using a 20-bp protospacer sequence within a guide RNA (gRNA)^2^. The 20-bp target site must be immediately upstream of a NRG sequence referred to as the protospacer adjacent motif (PAM)^3^ with an NGG site conferring increased cutting efficiency compared to NAG^4^. Given the potential number of matches and mismatches for a 20-bp sequence in large genomes along with reports of off-target Cas9 mutagenesis in cultured cells^5^, concerns regarding off-target Cas9 activity resulting in unintended genome modifications remains. In response, numerous methods have been developed to mitigate and detect purported off-target effects of Cas9 activity, such as the use of high fidelity Cas9 variants and gRNA modifications^6–10^, and unbiased molecular approaches to assess Cas9 off-target cutting: BLESS^11^, CIRCLE-seq^12^, Digenome-seq^13^, GUIDE-seq^14^, and SITE-seq^15^. However, these detection methods are often difficult to implement in large-scale animal production scenarios where the large number of embryos that would be required to implement these assays are ethically prohibitive in the absence of compelling arguments for their need. Further, the extent of reported Cas9-specific off-target mutagenesis varies across studies, ranging from almost undetectable to moderate when reagents are delivered directly to mouse zygotes^16–18^. These studies typically involve whole-genome sequencing for a limited number of gRNA targets with trios (parental-progeny) or intercrosses of inbred strains; and, thus diminish our ability to generalize these findings or interpret off-target events in the context of natural variation.

In order to assess the risk of unintentional off-target mutations when using Cas9 to create gene edited mouse lines, we compared whole-genome sequencing for 28 wild-type mice with 50 founder animals from the same C57BL/6N isogenic background generated using protocols and guides designed to minimize off-target risk (Supplemental Table S1). All CRISPR/Cas9-derived knock-out mice were generated using a deletion approach to ablate critical exon(s) using 2, 3 or 4 gRNA’s per target gene. Collectively, the founders represent 163 different guide RNA’s produced across four KOMP^2^ centers (Figure 1). In an attempt to replicate typical procedures used in mouse genetic engineering facilities, control samples were randomly selected from the production center’s wildtype C57BL/6N stud male colony used for embryo production or from the same embryo pool used to generate founders. Whole-genome sequencing was performed on individual samples to an average depth of ~35-40X coverage and processed for variant calling (Figure S1).

**Fig. 1.**
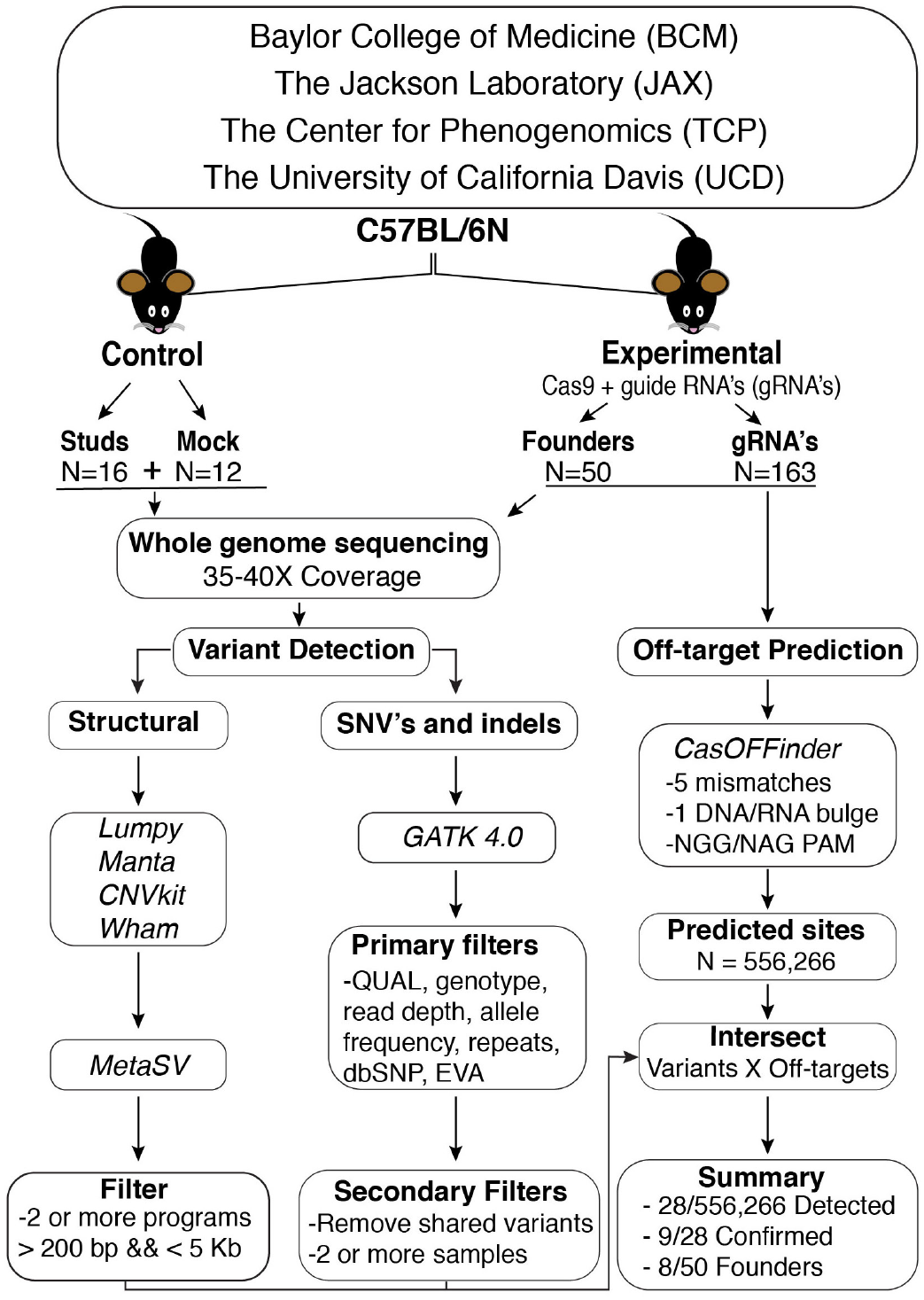
Multi-center analysis to assess off-target risk in CRISPR/Cas9 founder animals using whole genome sequencing (WGS). Genomic DNA from a subset of C57BL/B6N stud males used for embryo production or zygotes that were not treated with Cas9 was used as control DNA. Founders born from Cas9 gene editing experiments on zygotes from the stud males or from the same embryo pool comprised the experimental group. Each founder animal was created using a multi-guide strategy to delete a critical region. Founder animals were selected for WGS analysis after confirmation of germline transmission of the expected deletion. The whole-genome sequence analysis pipeline detected single nucleotide variants and small indels as well as potential structural variants. Potential off-target sites were predicted using CasOFFinder using permissive parameters and intersected with detected variants to identify putative off-target mutations.

For analysis, we first applied a primary filter to remove any variants found in dbSNP or the European Variant Archive (EVA) and a secondary filter to eliminate any variants shared between any two independent samples (Fig. 1). This process identified an average of 1,115 unique variants per control mouse and 1,034 variants per Cas9-treated mouse (Fig. 2a). Of these, ~756 single nucleotide variants (SNVs) and ~276 insertion/deletions (indels) were found per control sample and ~713 SNVs and ~322 indels per Cas9-treated mouse (Fig. 2a). No significant differences were observed between the total number or type of variants between control and Cas9-treated mice. Further, the position of variants did not measurably differ between groups, with most variants found within intergenic regions and introns and a reduced number observed in exons (Fig. 2b). The vast majority of the variants were heterozygous suggesting a large degree of diversity within each colony of animals (Fig. 2c). However, a large number of homozygous variants (n=2,876) were shared between at least 10 different samples, highlighting a number of C57BL/6N-specific variants currently not found in dbSNP or EVA (Fig. 2d).

**Fig. 2.**
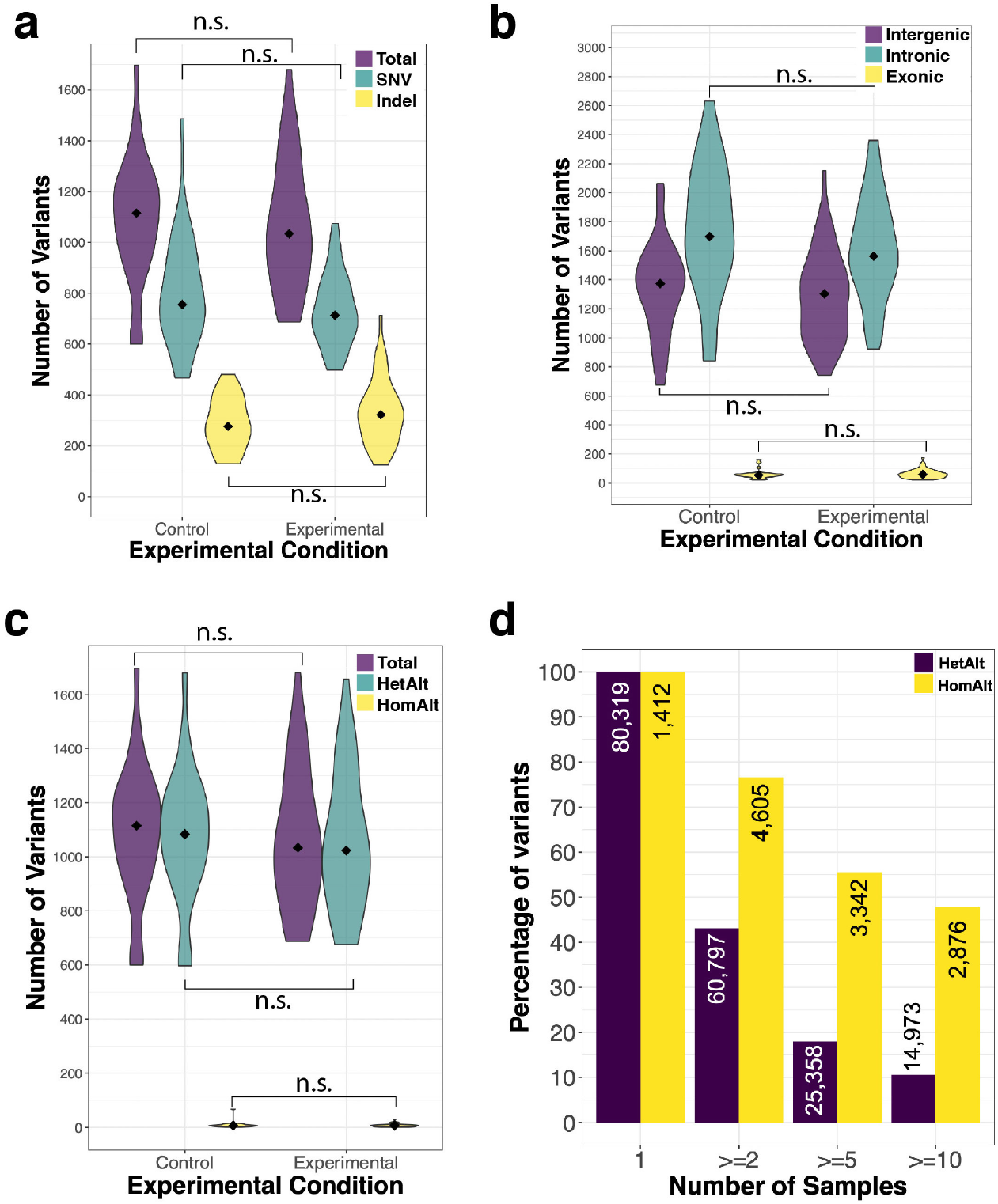
Summary of variants detected in control and Cas9 treated animals. **a.** Violin plots showing the total number of small variants identified within each experimental group. **b.** Distribution of variants throughout the genome relative to genic sequences. **c.** Zygosity of SNV and indel variants identified. **d.** Percentage of variants found in sample subsets. In violin plots, the median is denoted by a diamond. n.s. not significant (Wilcoxon test, α= 0.05).

Our variant calling pipeline successfully identified the expected exon deletion in 49/50 samples (98%). The single missed deletion corresponding to *Rasgef1a* was successfully identified by lumpy^19^ and manta^20^, but was later filtered out by manta due to low quality and thus failed to meet our threshold of being independently called by two or more programs (Supplemental Figure S2). Given the high concordance between WGS data and mutation detection, we set out to determine the extent to which unintended off-target effects were directly caused by spurious Cas9 activity. Cas-OFFinder^21^ was used to identify all predicted off-target sites associated with NGG/NAG PAM sequences and allowing for up to 5 mismatches with one DNA or RNA bulge compared to the on-target cut site. This resulted in 556,266 potential off-target regions in the genome for 163 tested gRNAs (Supplemental Table S2). From our WGS data, we detected off-target Cas9 activity at 0.005% (28/556,266) of predicted off-target sites associated with 18/50 samples tested resulting in less than one off-target hit per founder animal (Supplemental Table S3). Several genes (*Dmxl1*, *Fsd1l*, *Irf3, Plxnb1, Psma5, Ptp4a2*, and *Tmem171*) had more than one off-target event in the founder animal identified in the WGS data; however, three out of four variants called for *Tmem171* were SNVs associated with the same guide and were found in close proximity to one another within a region appearing to be highly polymorphic (Supplemental Figure S3q and Supplemental Table S3). Notably, the majority of pipeline predicted off-target sites were found in intergenic regions (Figure 3a) and there was a clear distinction between class of variant and PAM sequences with NAG primarily associated with structural variant calls and NGG with small indels (Figure 3b). WGS detected off-target activity was mostly associated with guides containing mismatches at distal PAM locations more tolerant to mutagenesis, consistent with the reported mechanisms of off-target activity^4^. However, some off-target activity was also noted even with mismatches at PAM proximal sites (Fig. 3c,d). All indels predicted to be the result of Cas9 off-target cutting were independently validated using Sanger sequencing (Supplemental Table S3). Despite previous reports of structural variants resulting from Cas9 activity^22^, we were only able to confirm small indels in close proximity to an NGG PAM (Supplemental Table S3 and Figure S3). These findings indicate that Cas9 off-target activity is predictable and can be minimized with careful guide selection.

**Fig. 3.**
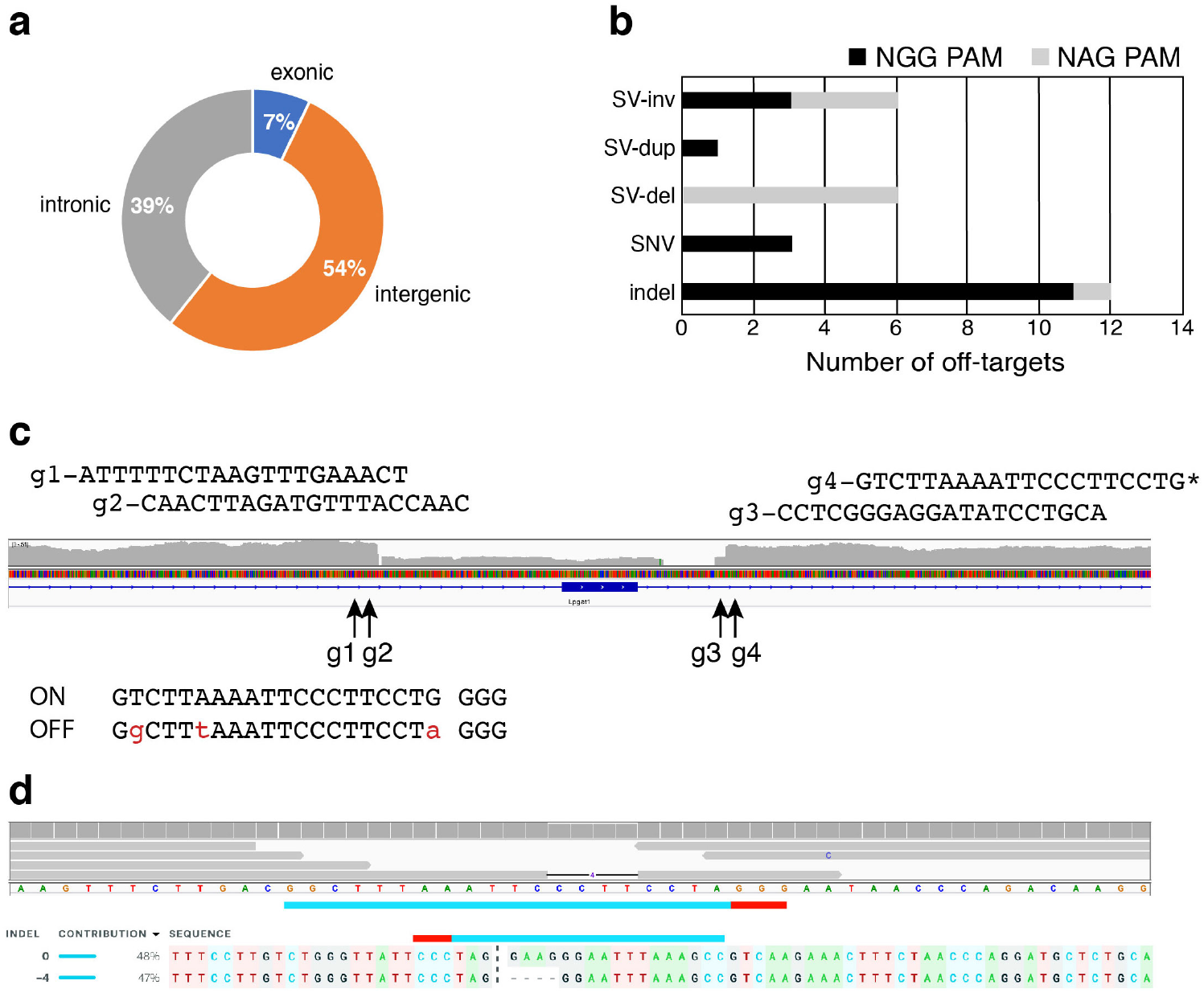
Detection of off-target Cas9 activity in whole genome sequencing data. **a.** Position of detected off-targets (N=28) relative to gene with percentage for each shown in doughnut plot. **b.** Classification of type of off-target mutation and associated PAM sequence of off-target guide. **c.** On target identification of exon deletion at *Lpgat1* generated using a four-guide design strategy. Off-target cutting was detected associated with guide sequence, g4. Mismatch sites are shown in red lowercase letters. **d.** Primary sequence data used to identify off-target site and Sanger Sequence validation from founder animal DNA confirming 4-bp deletion. ICE analysis shown below predicts a heterozygous allele frequency (https://www.synthego.com/products/bioinformatics/crispr-analysis). Abbreviations: SV, structural variant; inv, inversion; dup, duplication; del, deletion; SNV, single nucleotide variant; and indel, insertion/deletion.

In order to better understand the frequency of underlying genetic heterogeneity in inbred mice relative to the risk of Cas9 off-target activity, we analyzed the variants identified across all of the samples. We found the largest contributing factor was differences between the two C57BL/6N substrains used in this study (Fig. 4). Further, we did not observe an increase in the number of variants or segregation of Cas9 treated animals when compared to wild-type controls. These findings indicate that the diversity between any two individuals of the same substrain is greater than may be introduced by potential Cas9 off-target activity when using appropriately selected guides.

**Fig 4.**
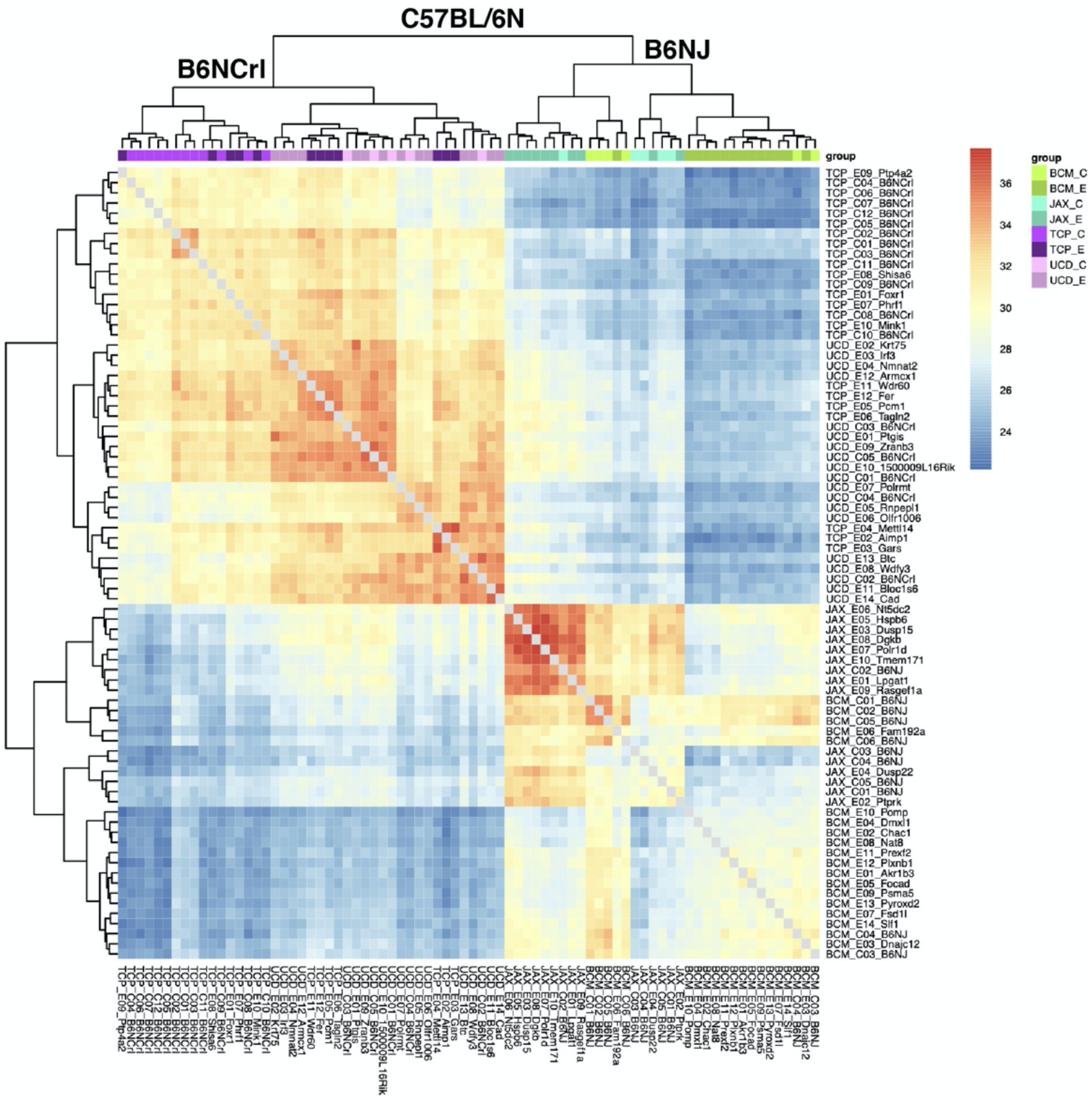
Genetic heterogeneity observed in individual mice of the same isogenic background. Heatmap shows the percentage of common SNP variants and highlights two major clusters defined by animal production center and mouse substrain used for genetic modification. C57BL/6NCrl (NCrl) mice were utilized by The Centre for Phenogenomics (TCP) and University of California, Davis (UCD); while C57BL/6NJ (NJ) mice were used by The Jackson Laboratory (JAX) and Baylor College of Medicine (BCM). Sample names are shown on the bottom and right side of figure using the production center abbreviation followed by target gene or substrain background and designation of treatment group either control (C) or experimental (E). Both treatment groups were interspersed with each other consistent with no statistical difference observed between control and experimental mice.

In this study, we set out to determine how frequently Cas9 off-target editing events occur in founder animals when applying well-defined design principles. While we did computationally detect unintentional Cas9-mediated off-target activity in 36% (18/50) of our lines, only 9/28 (32%) off-target events, associated with 16% (8/50) lines, were confirmed. Further, these off-target mutations were not linked to the intended target genes of interest. This outcome enables segregation of the off-target mutations via backcrossing that normally occurs during breeding for line expansion or during intercross of heterozygous mice to produce experimental and control cohorts, thereby controlling for both naturally occurring and Cas9-meditated off-target mutations. Furthermore, large-scale off-target structural variants were not found, and the frequency of off-target mutations occurred far less often compared to naturally occurring sequence variation found between any two mice of the same substrain.

Here, we provide WGS data for a far greater number of gRNA target sites than in previous reports^16–18^. Our study design captured the intrinsic heterogeneity present in a given inbred strain as they are typically maintained at a vendor or an accredited mouse breeding facility. Operationally, it is not feasible to obtain parental information for the pool of zygotes generated from multiple breeding pairs necessary to perform gene editing experiments at large-scale or even in typical production workflows in most core facilities. Although inbred strains are assumed to be isogenic, the spontaneous, *de novo* mutation rate in mice has been estimated to be between ~50-100 SNVs and 3-4 indels per generation^23,24^. Therefore, generations of even carefully maintained colonies will accumulate a significant number of variants due to genetic drift as they are expanded from foundation stocks. Our results clearly show that the rate of Cas9-induced off-target mutagenesis in a carefully designed mouse production experiment is trivial relative to the overall genetic heterogeneity observed in carefully maintained inbred mouse colonies. With this in mind, we strongly recommend the selection of appropriate genetic controls, such as littermates or inbred animals taken at random from the colony used for backcross breeding to mutants, for comparison when assessing gene-phenotype relationships in modified animals. Further, it is important to note that backcrossing or outcrossing mice introduces significantly more variation than the use of Cas9 and the appropriate control animals for most genetic experiments are littermate or line mate wild-type mice.

In summary, these data indicate that the risk of Cas9 cutting at predicted off-target sites is significantly lower than the natural genetic variation introduced into the genomes of inbred mice through natural mating. It is important to note that for highly specific gene editing experiments requiring the use of guides that may have increased off-target risk, it is advised to check for these events in both founder animals and in the N1 generation, particularly if the predicted mutations are genetically linked to the target or occur in an exon or functional sequence element.

## Materials & Methods

### Allele design and guide selection

For multi-exon genes, a critical region (one or more exons) was identified as shared among all annotated full-length transcripts whose removal was predicted to result in a frame-shift mutation and introduction of premature stop codon resulting in nonsense mediated decay of mRNA when deleted. gRNA sequences flanking the critical region were chosen using the constraints that they had no off-target sites with less than three mismatches adjacent to an NGG PAM. Guides were prioritized to minimize off-target risk and maximize predicted on-target cutting efficiency using prediction algorithms including CRISPRtools^25^, CRISPR MIT, CHOPCHOP^26^, CRISPOR^27^, and WGE^28^. Guide information is summarized in Supplemental Table S2.

### Animals

All experiments were performed on C57BL/B6N mice obtained from either The Jackson Laboratory (C57BL/6NJ; stock #5304) or Charles River (C57BL/6NCrl; strain code 027). All animals were maintained in accordance with institutional policies governing the ethical care and use of animals in research under approved protocols. All procedures involving animals at The Centre for Phenogenomics (TCP) were performed in compliance with the Animals for Research Act of Ontario and the Guidelines of the Canadian Council on Animal Care under Animal Use Protocols 0008, 0084 and 0275 reviewed and approved by the TCP’s Animal Care Committee. All animal use at Baylor College of Medicine, The Jackson Laboratory and UC Davis were done in accordance with the Animal Welfare Act and the AVMA Guidelines on Euthanasia, in compliance with the ILAR Guide for Care and Use of Laboratory Animals, and with prior approval from their respective institutional animal care and use committees (IACUC).

### Cas9 and guide RNA delivery to zygotes

Gene editing was performed by either microinjection or electroporation of Cas9 mRNA and gRNA or Cas9 ribonucleoprotein complexes (RNP), respectively. Electroporation and microinjection experiments were conducted essentially as previously described^29–31^.

### Whole genome sequencing

Genomic DNA for founder and control samples was extracted from spleens, tail tips or ear punches using phenol:chloroform or kit according to manufacturer’s suggestions. DNA was quantified using fluorescence-based detection on a Qubit (Thermofisher). Whole genome sequencing libraries were prepared at The Centre for Applied Genomics (The Hospital of Sick Children, Toronto, Ontario) following standard practices. Briefly, 700 ng of genomic DNA was sheared to an average size of 400 bp using a Covaris LE220 and was used as input to generate a whole-genome library using the TruSeq PCR-free kit (Illumina). The resulting DNA libraries were sequenced on Illumina Hi-seq X instrument to generate 2×150bp paired-end reads.

Sequence data associated with this study are deposited to the NCBI Sequence Read Archive (SRA) under accession number PRJNA687003.

### NGS data analysis

Sequence read quality was assessed using FastqC (https://www.bioinformatics.babraham.ac.uk/projects/fastqc/) and fastQ Screen^32^, reads were processed through the bcbio pipeline (https://github.com/bcbio/bcbio-nextgen, ver. 1.1.0) for all steps from alignment to variant calling. Briefly, reads were aligned to mouse genome assembly (GRC Build 38/mm10) using bwa-mem^33^ resulting in ~35-40X genome coverage for each sample. GATK 4.0 (Genome Analysis Toolkit)^34,35^ was used to call variants with the default parameters of the pipeline. The resulting variant call format (VCF) files were filtered to retain variants with QUAL>30, DP>9, GQ>30 and AF>0.1 using bcftools (ver. 1.6)^36^. Repetitive intervals were padded by two base pairs on either side to improve filtering due to indel variants that overlap boundaries of the repetitive intervals. Subsequently, variants were filtered using the central repository for mouse, European Variation Archive (EVA, https://www.ebi.ac.uk/eva), and dbSNP. A non-redundant set of variants was obtained by merging EVA files (GRCm38.p2, GRCm38p3, and GRCm38p4) that were then applied to filter out any common variants present in the 78 VCF files. Using a custom python script, heterozygous variants with a ratio of alternative alleles to total number of alleles less than 0.2 were excluded. Callable intervals were defined by bcbio pipeline for each sample based on the corresponding bam file. Since the number of variants for each sample, which is the main parameter in our analysis, can be influenced by the extent of callable intervals, we set to limit the primary-filtered VCF files to the intersection of all samples callable intervals to avoid any potential bias. Callable interval filtering was performed using a combination of custom-made scripts and bedtools *multiinter* tool (ver. 2.27.1)^37^. The intersection of callable intervals common to all 78 samples was used to filter the variants outside these intervals. A final set of variants was determined by applying a secondary filter to eliminate any variants observed in two or more independent samples using the bcbio-variation-recall ensemble software (https://github.com/bcbio/bcbio.variation.recall, ver. 0.1.7). bcftools *isec* was then applied to filter out the ensemble file variants from each sample’s VCF file to create the secondary-filtered VCF files. Structural variants (SV’s) were called by lumpy^19^, manta^20^, CNVkit^38^, and Wham^39^ followed by Metasv^40^. After removing ./. and 0/0 genotypes, SVs considered for final analysis had to be called by 2 or more methods and have at least 3 reads supporting with a variant length greater than 200 bp but less than 5 kb. SnpEff (ver. 4.3t)^41^ was used to divide each sample’s secondary filtered variants into intergenic, exonic, and intronic regions. The downstream and upstream variants were included in the intergenic category. For the comparison of the average means between experimental and control groups, the “compare_means” method of R package ggpubr (https://rpkgs.datanovia.com/ggpubr/) was used which provided the Wilcoxon test followed by Bonferroni correction. The bcftools *isec* tool was used to find the number of overlapping and unique variants between any given three primary-filtered VCF files. The primary-filtered VCF files (excluding the dbSNP150, EVA, and callableIntervals filters mentioned above) were submitted to the European Variation Archive (EVA) database (https://www.ebi.ac.uk/eva/). Predicted off-targets were identified using Cas-OFFinder^21^ and visually assessed using IGV (Integrated Genome Viewer). Heatmap dendrogram was created by providing the percentage of common variants between any two samples (using “bcftools isec” tool and python scripts) as input to the pheatmap R package (https://cran.r-project.org/web/packages/pheatmap/).

### Off-target validation

PCR products were generated from founder and control DNA samples using specific primers surrounding the region of interest. PCR products were submitted for Sanger Sequencing and analyzed using ICE (https://www.synthego.com/products/bioinformatics/crispr-analysis).

## Acknowledgements

We would like to thank The Centre for Applied Genomics for generating whole genome sequencing data. We also thank the model production staff at The Jackson Laboratory, The Centre for Phenogenomics, Baylor College of Medicine, and UC Davis Mouse Biology Program for their assistance generating the mouse mutants used for this study. Research reported in this work was supported by the NIH Common Fund, National Human Genome Research Institute UM1 HG006348 (J.D.H. and M.E.D.), and National Institutes of Health Office of the Director UM1 OD023221 (K.C.K.L.) and UM1 OD023222 (R.E.B., S.A.M. and J.K.W.) and Genome Canada and Ontario Genomics OGI-137 (L.M.J.N.). Additional support was provided by the Canadian Center for Computational Genomics (C3G), part of the Genome Technology Platform (GTP), funded by Genome Canada through Genome Québec and Ontario Genomics (A.R.). The content is solely the responsibility of the authors and does not necessarily represent the official views of the National Institutes of Health.

## Author Contributions

JDH, MED, KCKL, LMJN and SAM conceived the project and received funding; JDH, DGL, BW, JAW, LMJN, KAP, KCKL, and SAM produced Cas9 knockout mice; AR, KAP, NH and SK analyzed the data. DGL, LGL, BW, LMJN and KAP performed off-target validation experiments. KAP, SK and NH generated figures and drafted the manuscript. All authors provided feedback and edits to the manuscript.

## Competing Interests

The authors declare no competing interests.

## Notes

https://www.ncbi.nlm.nih.gov/sra

### Competing Interest Statement

The authors have declared no competing interest.

## References

1 Saha, K. et al. The NIH Somatic Cell Genome Editing program. Nature 592, 195–204, doi:10.1038/s41586-021-03191-1 (2021).

2 Doudna, J. A. & Charpentier, E. Genome editing. The new frontier of genome engineering with CRISPR-Cas9. Science 346, 1258096, doi:10.1126/science.1258096 (2014).

3 Jinek, M. et al. A programmable dual-RNA-guided DNA endonuclease in adaptive bacterial immunity. Science 337, 816–821, doi:10.1126/science.1225829 (2012).

4 Hsu, P. D. et al. DNA targeting specificity of RNA-guided Cas9 nucleases. Nat Biotechnol 31, 827–832, doi:10.1038/nbt.2647 (2013).

5 Fu, Y. et al. High-frequency off-target mutagenesis induced by CRISPR-Cas nucleases in human cells. Nat Biotechnol 31, 822–826, doi:10.1038/nbt.2623 (2013).

6 Chen, J. S. et al. Enhanced proofreading governs CRISPR-Cas9 targeting accuracy. Nature 550, 407–410, doi:10.1038/nature24268 (2017).

7 Fu, Y., Sander, J. D., Reyon, D., Cascio, V. M. & Joung, J. K. Improving CRISPR-Cas nuclease specificity using truncated guide RNAs. Nat Biotechnol 32, 279–284, doi:10.1038/nbt.2808 (2014).

8 Kleinstiver, B. P. et al. High-fidelity CRISPR-Cas9 nucleases with no detectable genome-wide off-target effects. Nature 529, 490–495, doi:10.1038/nature16526 (2016).

9 Slaymaker, I. M. et al. Rationally engineered Cas9 nucleases with improved specificity. Science 351, 84–88, doi:10.1126/science.aad5227 (2016).

10 Vakulskas, C. A. et al. A high-fidelity Cas9 mutant delivered as a ribonucleoprotein complex enables efficient gene editing in human hematopoietic stem and progenitor cells. Nat Med 24, 1216–1224, doi:10.1038/s41591-018-0137-0 (2018).

11 Crosetto, N. et al. Nucleotide-resolution DNA double-strand break mapping by next-generation sequencing. Nat Methods 10, 361–365, doi:10.1038/nmeth.2408 (2013).

12 Tsai, S. Q. et al. CIRCLE-seq: a highly sensitive in vitro screen for genome-wide CRISPR-Cas9 nuclease off-targets. Nat Methods 14, 607–614, doi:10.1038/nmeth.4278 (2017).

13 Kim, D. et al. Digenome-seq: genome-wide profiling of CRISPR-Cas9 off-target effects in human cells. Nat Methods 12, 237–243, 231 p following 243, doi:10.1038/nmeth.3284 (2015).

14 Tsai, S. Q. et al. GUIDE-seq enables genome-wide profiling of off-target cleavage by CRISPR-Cas nucleases. Nat Biotechnol 33, 187–197, doi:10.1038/nbt.3117 (2015).

15 Cameron, P. et al. Mapping the genomic landscape of CRISPR-Cas9 cleavage. Nat Methods 14, 600–606, doi:10.1038/nmeth.4284 (2017).

16 Anderson, K. R. et al. CRISPR off-target analysis in genetically engineered rats and mice. Nat Methods 15, 512–514, doi:10.1038/s41592-018-0011-5 (2018).

17 Iyer, V. et al. No unexpected CRISPR-Cas9 off-target activity revealed by trio sequencing of gene-edited mice. PLoS Genet 14, e1007503, doi:10.1371/journal.pgen.1007503 (2018).

18 Willi, M., Smith, H. E., Wang, C., Liu, C. & Hennighausen, L. Mutation frequency is not increased in CRISPR-Cas9-edited mice. Nat Methods 15, 756–758, doi:10.1038/s41592-018-0148-2 (2018).

19 Layer, R. M., Chiang, C., Quinlan, A. R. & Hall, I. M. LUMPY: a probabilistic framework for structural variant discovery. Genome Biol 15, R84, doi:10.1186/gb-2014-15-6-r84 (2014).

20 Chen, X. et al. Manta: rapid detection of structural variants and indels for germline and cancer sequencing applications. Bioinformatics 32, 1220–1222, doi:10.1093/bioinformatics/btv710 (2016).

21 Bae, S., Park, J. & Kim, J. S. Cas-OFFinder: a fast and versatile algorithm that searches for potential off-target sites of Cas9 RNA-guided endonucleases. Bioinformatics 30, 1473–1475, doi:10.1093/bioinformatics/btu048 (2014).

22 Shin, H. Y. et al. CRISPR/Cas9 targeting events cause complex deletions and insertions at 17 sites in the mouse genome. Nat Commun 8, 15464, doi:10.1038/ncomms15464 (2017).

23 Lindsay, S. J., Rahbari, R., Kaplanis, J., Keane, T. & Hurles, M. E. Similarities and differences in patterns of germline mutation between mice and humans. Nat Commun 10, 4053, doi:10.1038/s41467-019-12023-w (2019).

24 Uchimura, A. et al. Germline mutation rates and the long-term phenotypic effects of mutation accumulation in wild-type laboratory mice and mutator mice. Genome Res 25, 1125–1134, doi:10.1101/gr.186148.114 (2015).

25 Peterson, K. A. et al. CRISPRtools: a flexible computational platform for performing CRISPR/Cas9 experiments in the mouse. Mamm Genome 28, 283–290, doi:10.1007/s00335-017-9681-z (2017).

26 Labun, K. et al. CHOPCHOP v3: expanding the CRISPR web toolbox beyond genome editing. Nucleic Acids Res 47, W171–W174, doi:10.1093/nar/gkz365 (2019).

27 Concordet, J. P. & Haeussler, M. CRISPOR: intuitive guide selection for CRISPR/Cas9 genome editing experiments and screens. Nucleic Acids Res 46, W242–W245, doi:10.1093/nar/gky354 (2018).

28 Hodgkins, A. et al. WGE: a CRISPR database for genome engineering. Bioinformatics 31, 3078–3080, doi:10.1093/bioinformatics/btv308 (2015).

29 Modzelewski, A. J. et al. Efficient mouse genome engineering by CRISPR-EZ technology. Nat Protoc 13, 1253–1274, doi:10.1038/nprot.2018.012 (2018).

30 Wang, W. et al. Delivery of Cas9 Protein into Mouse Zygotes through a Series of Electroporation Dramatically Increases the Efficiency of Model Creation. J Genet Genomics 43, 319–327, doi:10.1016/j.jgg.2016.02.004 (2016).

31 Gertsenstein, M. & Nutter, L. M. J. Production of knockout mouse lines with Cas9. Methods, doi:10.1016/j.ymeth.2021.01.005 (2021).

32 Wingett, S. W. & Andrews, S. FastQ Screen: A tool for multi-genome mapping and quality control. F1000Res 7, 1338, doi:10.12688/f1000research.15931.2 (2018).

33 Li, H. Aligning sequence reads, clone sequences and assembly contigs with BWA-MEM. arXiv:1303.3997v2 [q-bio.GN] (2013).

34 McKenna, A. et al. The Genome Analysis Toolkit: a MapReduce framework for analyzing next-generation DNA sequencing data. Genome Res 20, 1297–1303, doi:10.1101/gr.107524.110 (2010).

35 Van der Auwera, G. A. et al. From FastQ data to high confidence variant calls: the Genome Analysis Toolkit best practices pipeline. Curr Protoc Bioinformatics 43, 11 10 11–11 10 33, doi:10.1002/0471250953.bi1110s43 (2013).

36 Li, H. et al. The Sequence Alignment/Map format and SAMtools. Bioinformatics 25, 2078–2079, doi:10.1093/bioinformatics/btp352 (2009).

37 Quinlan, A. R. & Hall, I. M. BEDTools: a flexible suite of utilities for comparing genomic features. Bioinformatics 26, 841–842, doi:10.1093/bioinformatics/btq033 (2010).

38 Talevich, E., Shain, A. H., Botton, T. & Bastian, B. C. CNVkit: Genome-Wide Copy Number Detection and Visualization from Targeted DNA Sequencing. PLoS Comput Biol 12, e1004873, doi:10.1371/journal.pcbi.1004873 (2016).

39 Kronenberg, Z. N. et al. Wham: Identifying Structural Variants of Biological Consequence. PLoS Comput Biol 11, e1004572, doi:10.1371/journal.pcbi.1004572 (2015).

40 Mohiyuddin, M. et al. MetaSV: an accurate and integrative structural-variant caller for next generation sequencing. Bioinformatics 31, 2741–2744, doi:10.1093/bioinformatics/btv204 (2015).

41 Cingolani, P. et al. A program for annotating and predicting the effects of single nucleotide polymorphisms, SnpEff: SNPs in the genome of Drosophila melanogaster strain w1118; iso-2; iso-3. Fly (Austin) 6, 80–92, doi:10.4161/fly.19695 (2012).

