## Supplemental Material for "Whole genome analysis for 163 guide RNAs in Cas9 edited mice reveals minimal off-target activity"

**Supplemental Materials**

**Table S1.** Sample information for whole genome sequence analysis.

**Table S2**. Guide RNA information and predicted off-target sites.

**Table S3.** Variant calls for off-target sites detected in sequence analysis pipeline.

### Supplemental Figure S1

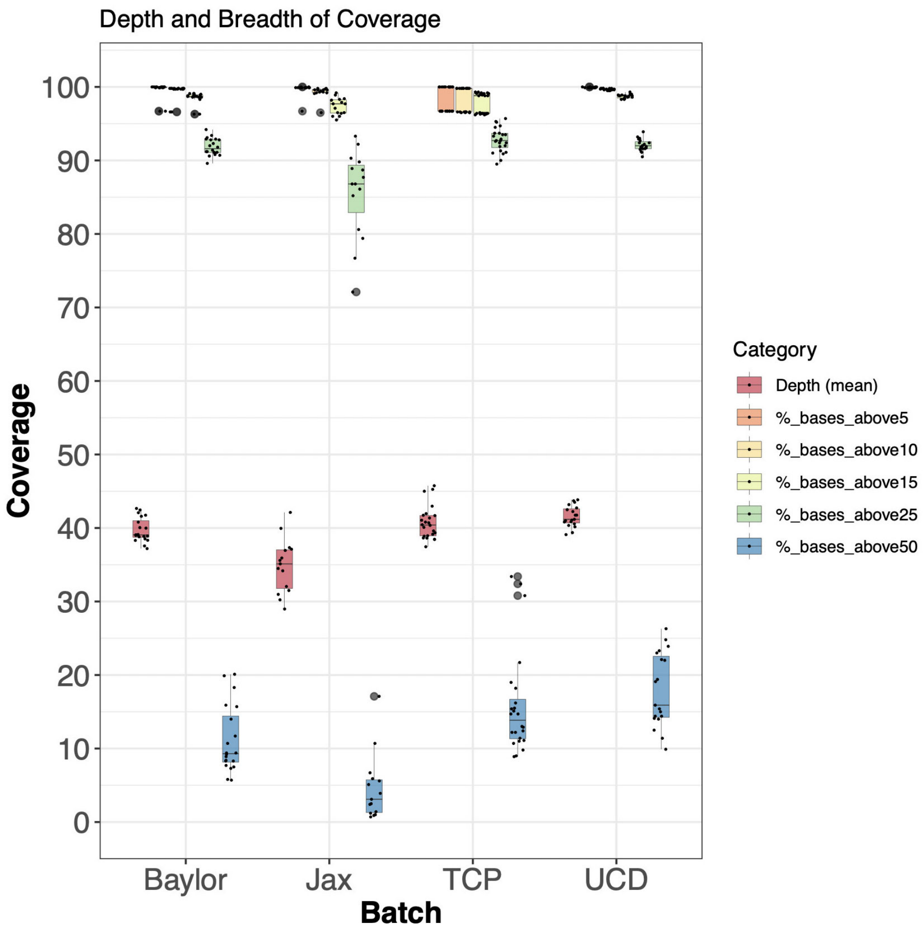


**Figure S1.** Summary of whole-genome sequencing data obtained from each KOMP2 production center showing base pair quality and depth.

### Supplemental Figure S2

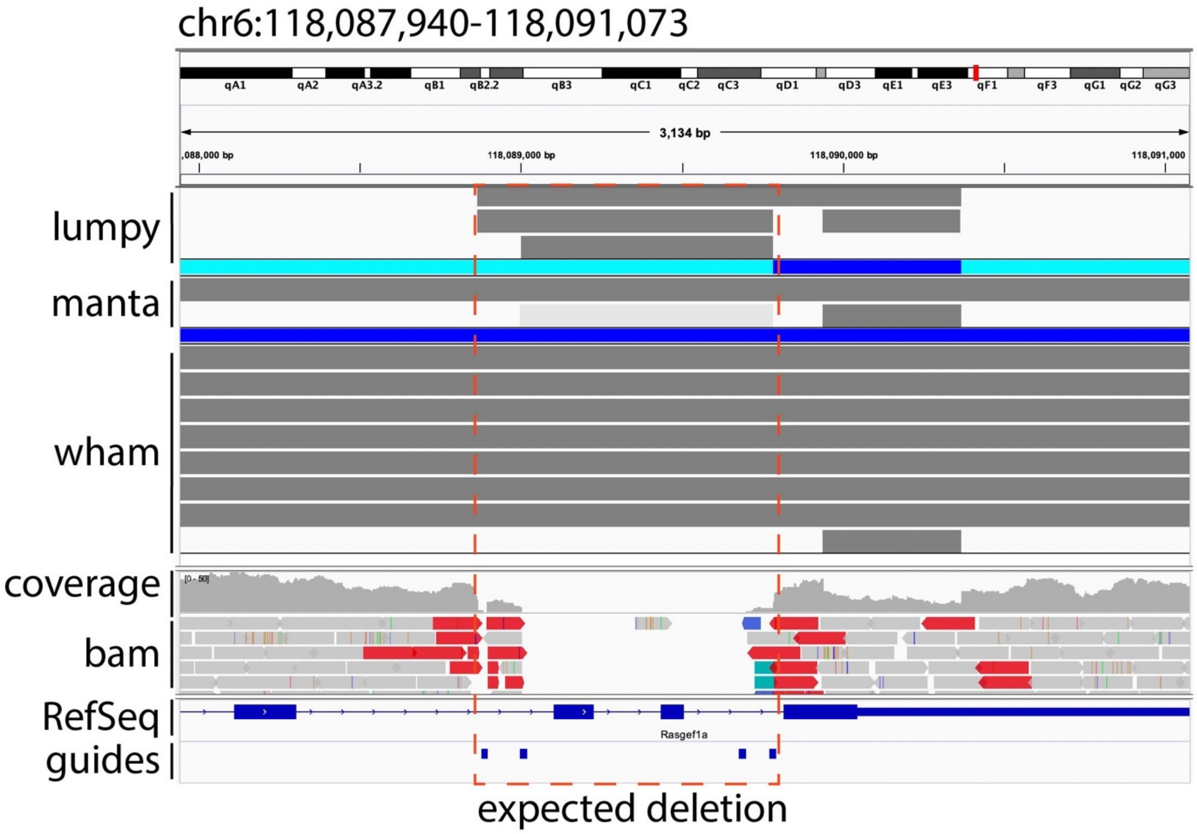


**Figure S2**. Comparison of structural variant callers for *Rasgef1a* deletion. The expected region is depicted with a red box and shows reduced sequence coverage consistent with Cas9 mediated deletion using a four-guide design strategy. The deletion was successfully called by lumpy and manta but filtered out by manta due to low confidence genotype filter indicted by light gray bar. This filtering resulted in the *Rasgef1a* deletion failing to meet the minimum threshold of being independently identified by at least two programs.

### Supplemental Figure S3

**a**

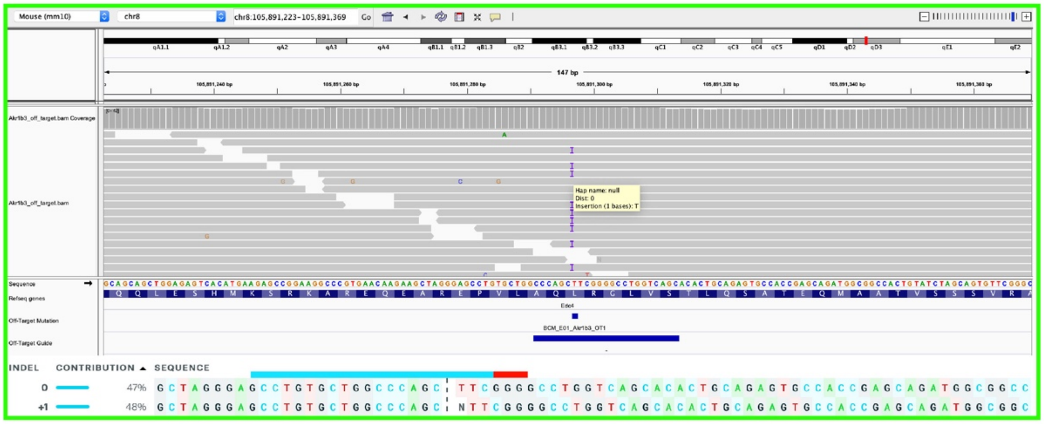

**b**

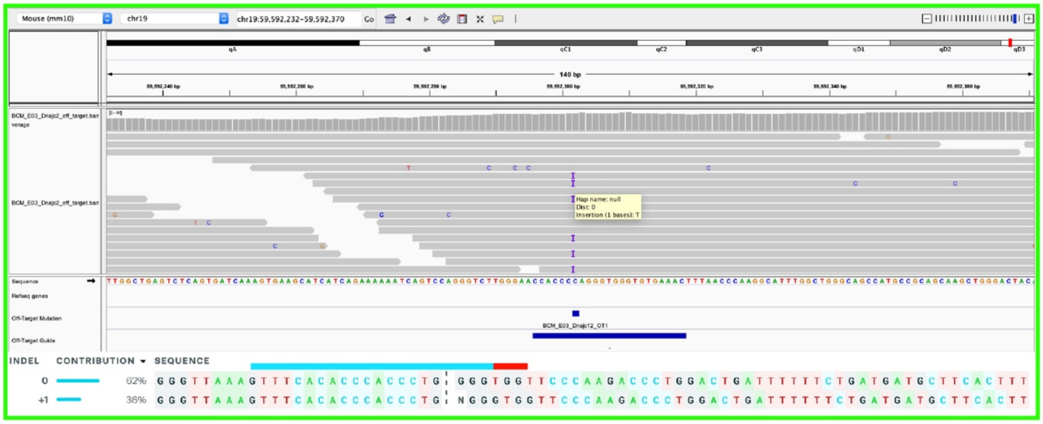

**c**

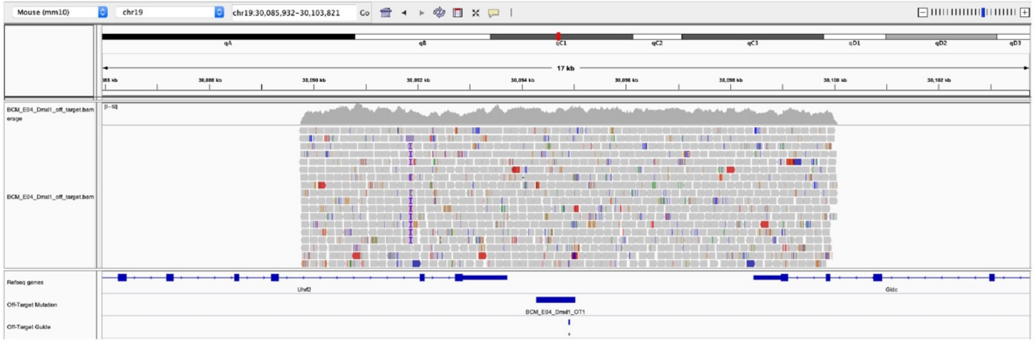

### Supplemental Figure S3, cont

**d**

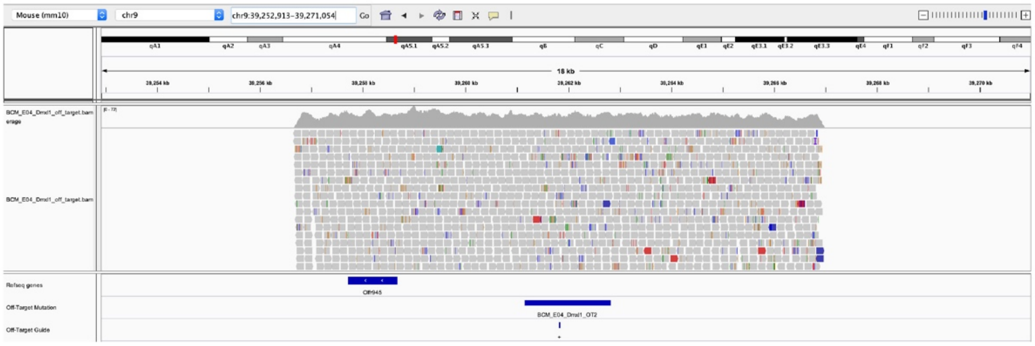


**e**

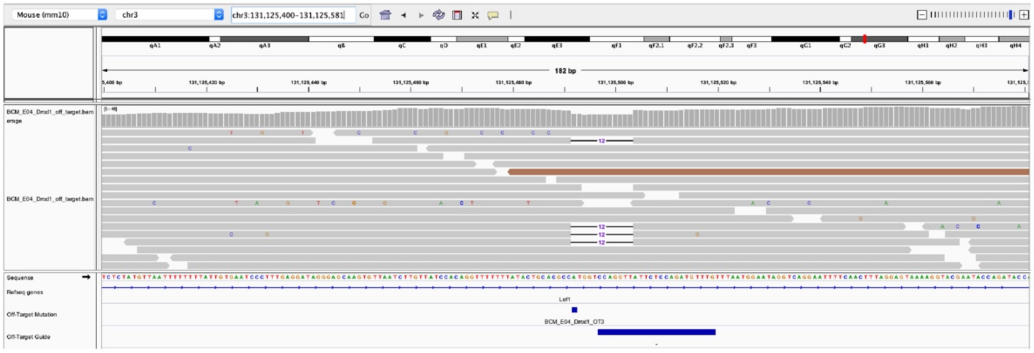


**f**

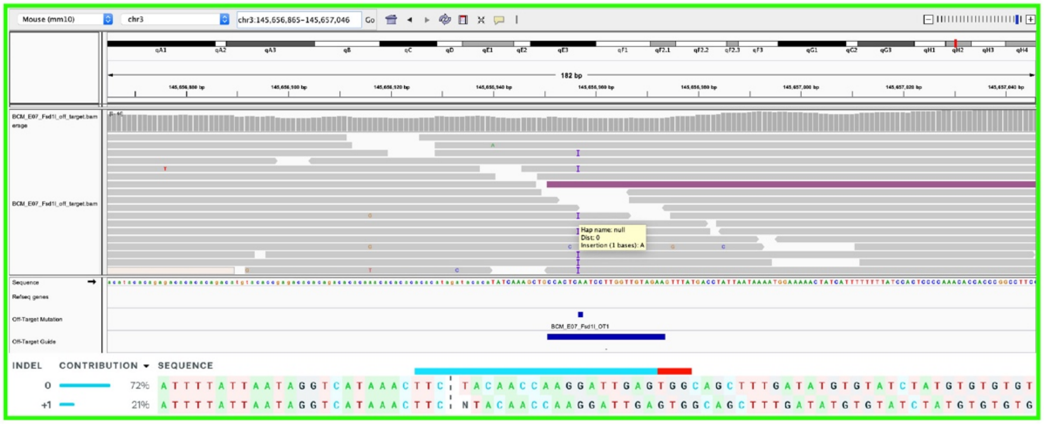

### Supplemental Figure S3,cont

**g**

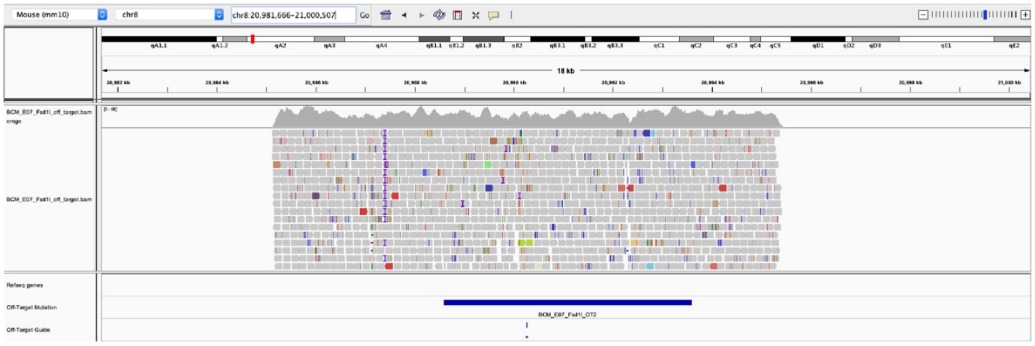


**h**

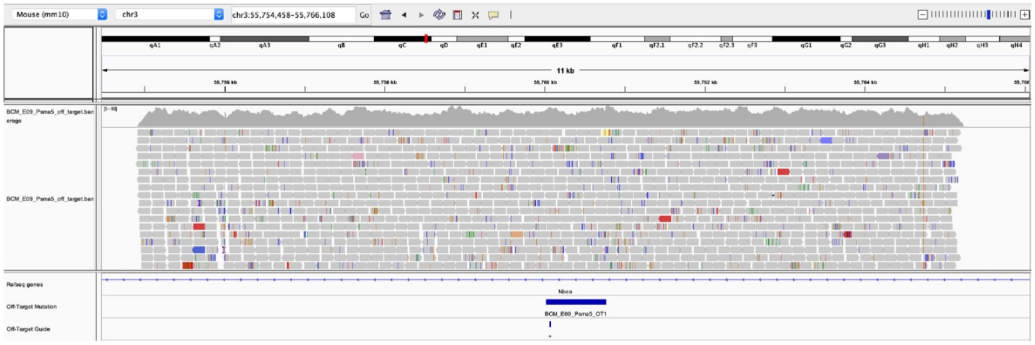


**i**

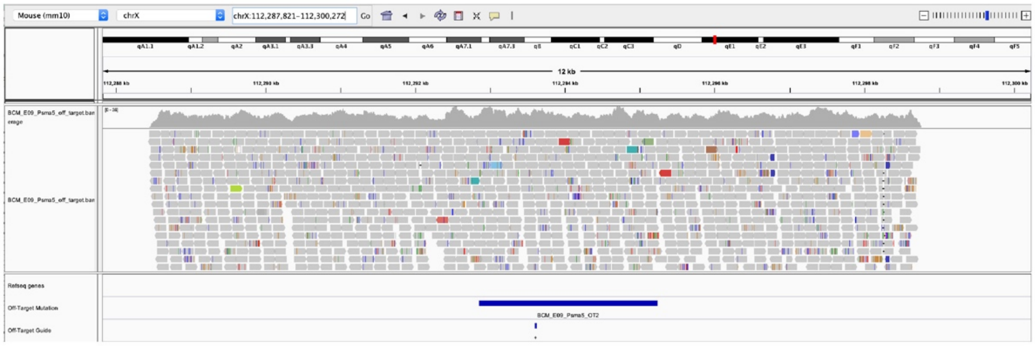

#### Supplemental Figure S3,cont

j

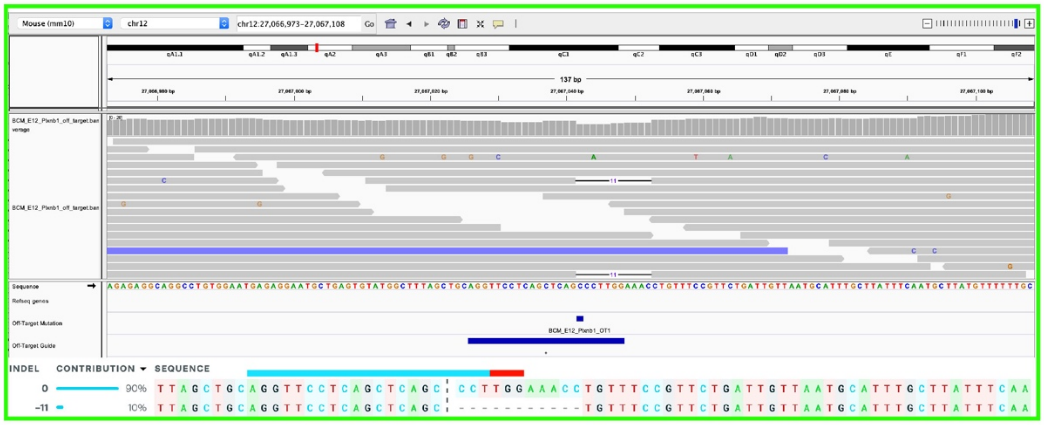

k

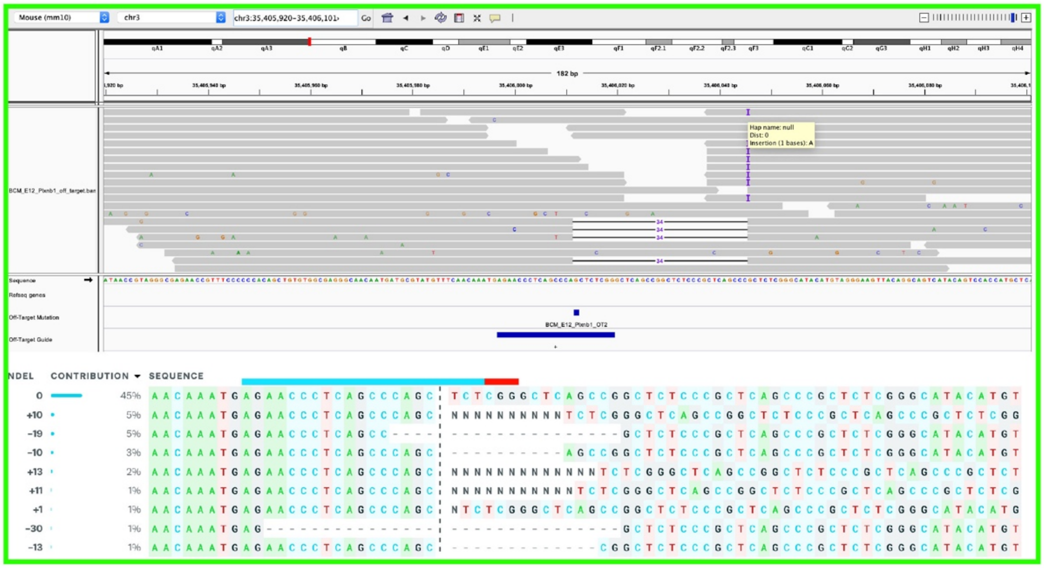

### Supplemental Figure S3,cont

l

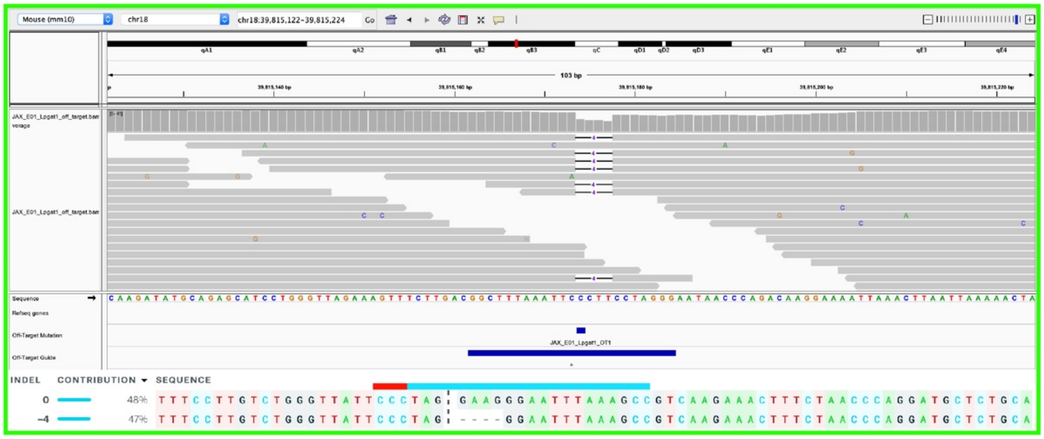


m

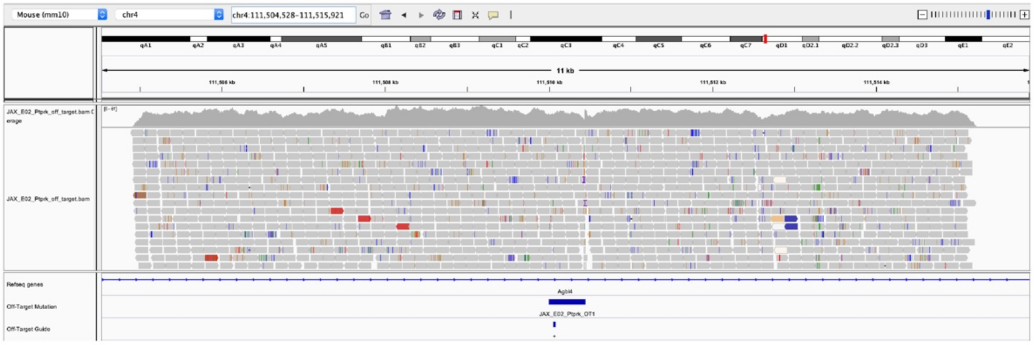


n

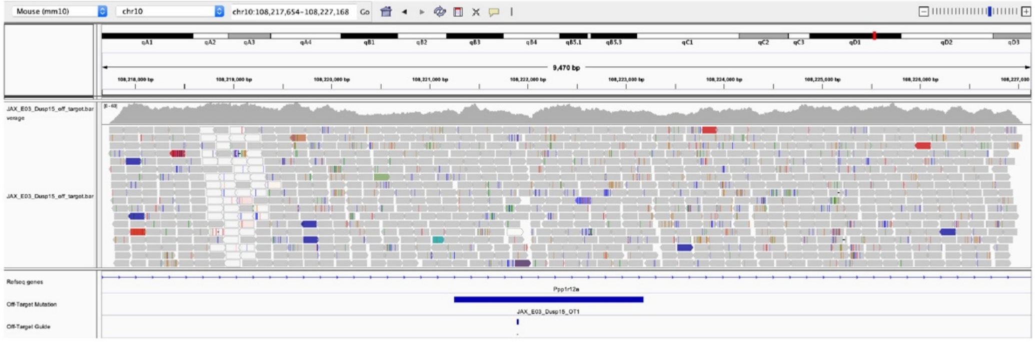

### Supplemental Figure S3,cont

**o**

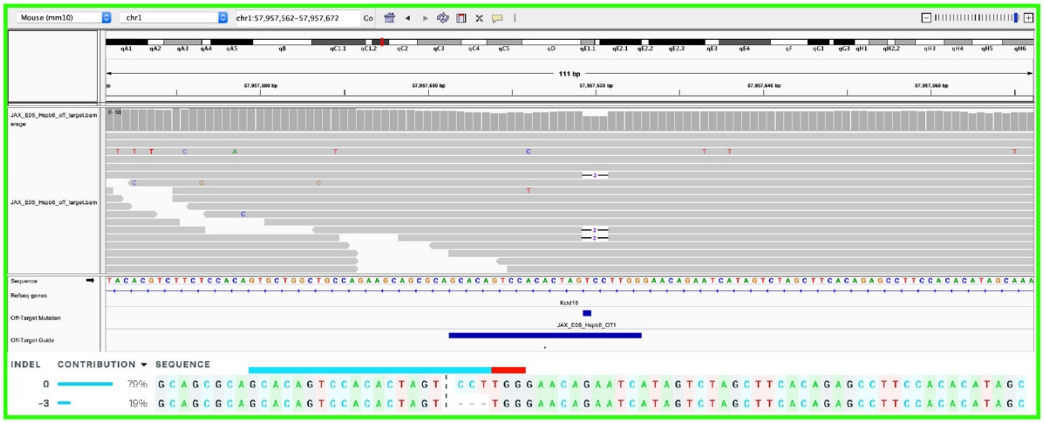


**p**

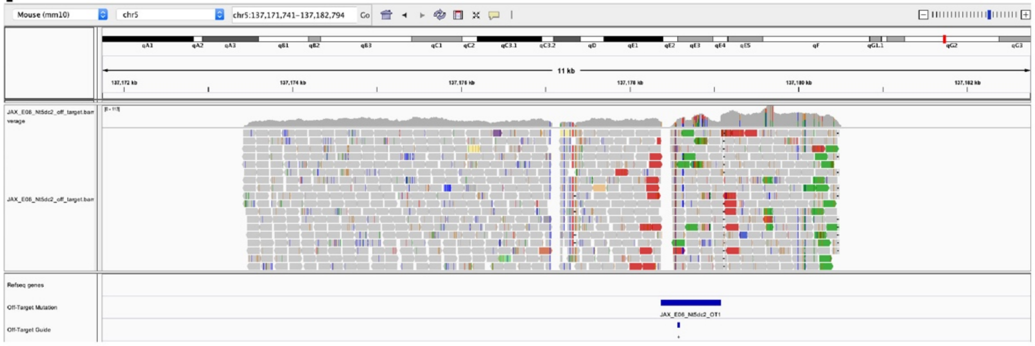


**q**

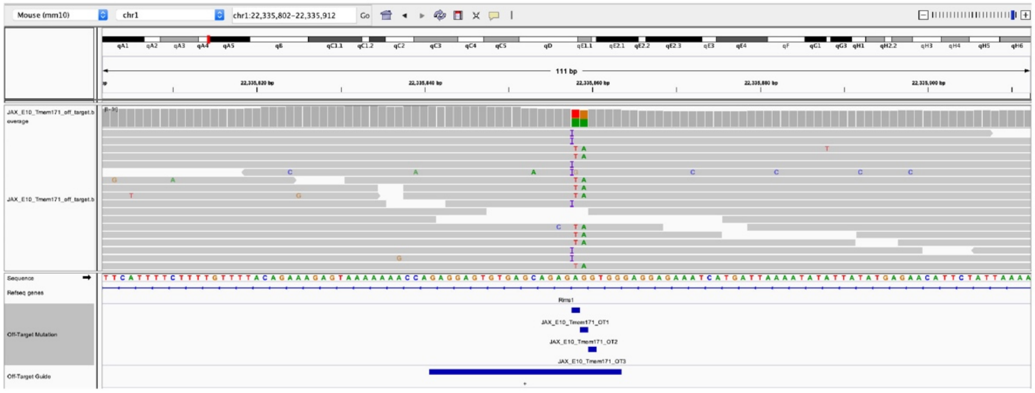

### Supplemental Figure S3,cont

**r**

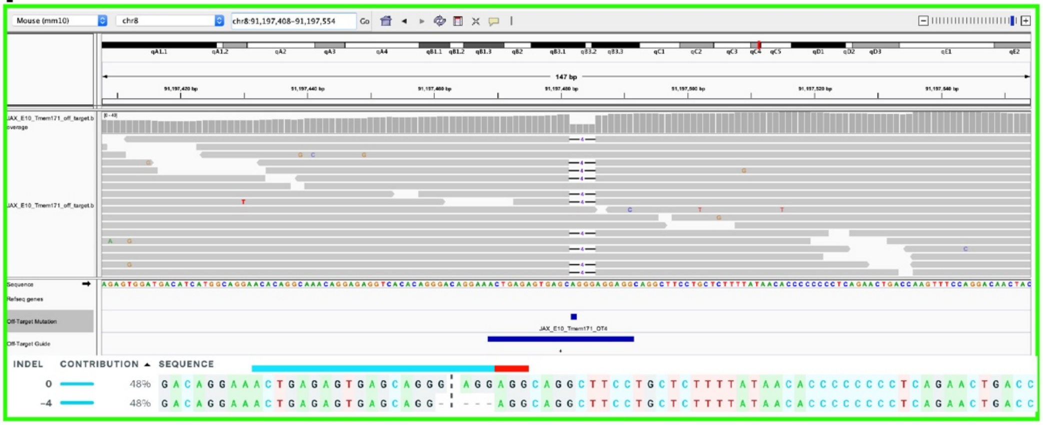


**s**

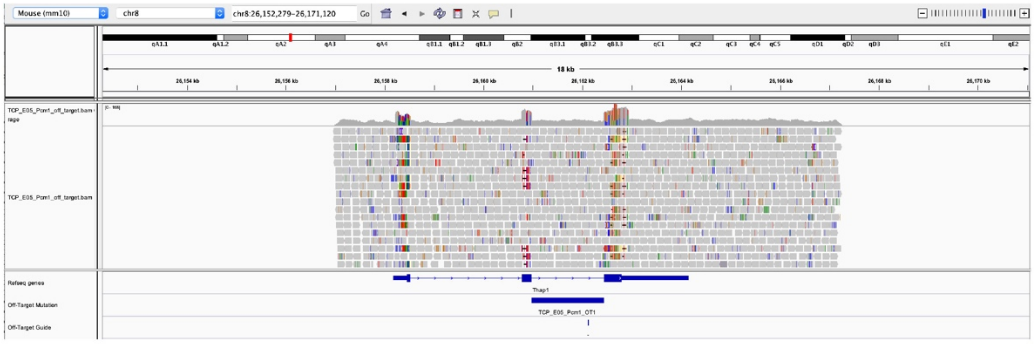


**t**

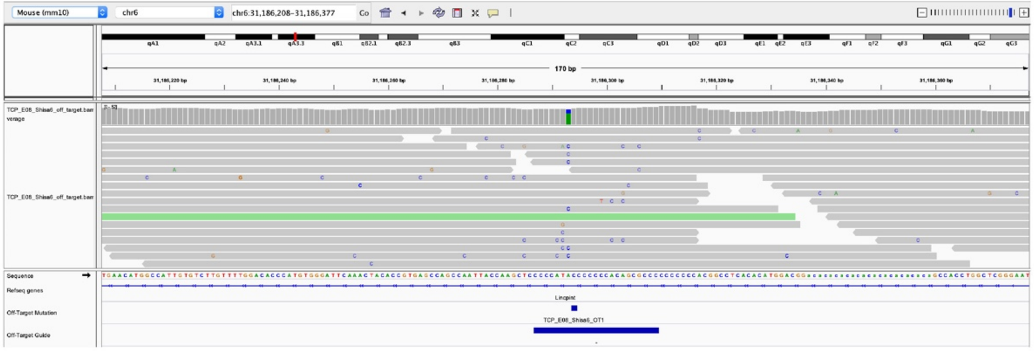

### Supplemental Figure S3,cont

**u**

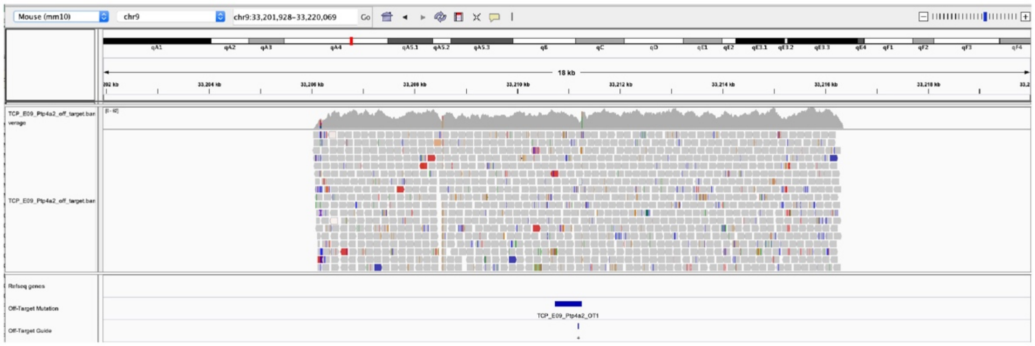


**v**

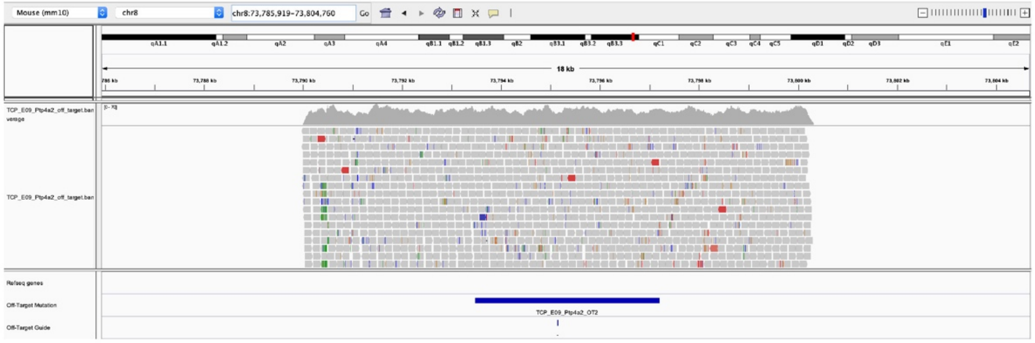


**w**

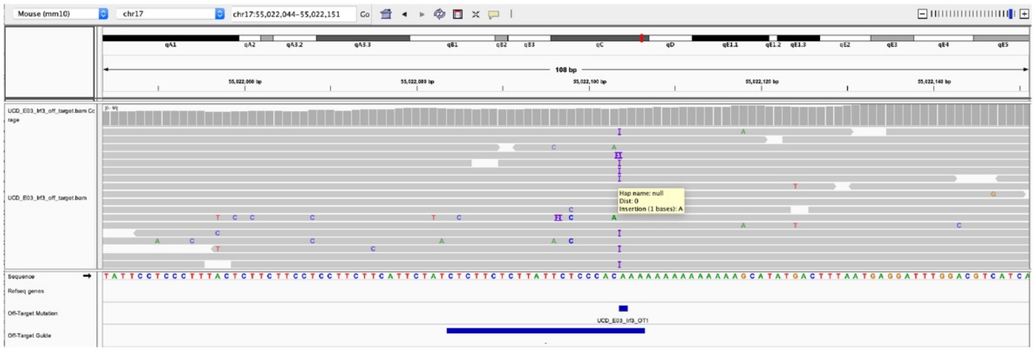

### Supplemental Figure S3,cont

x

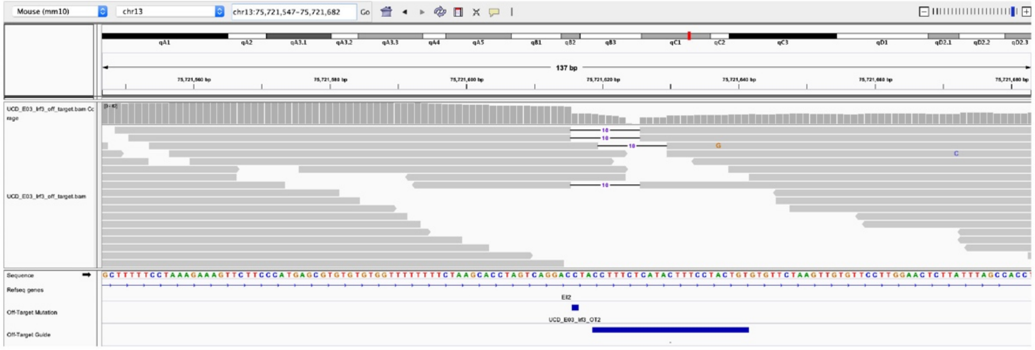

y

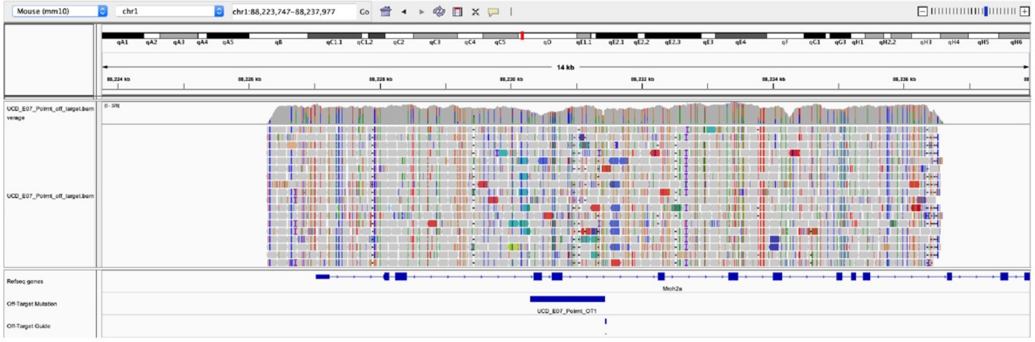

z

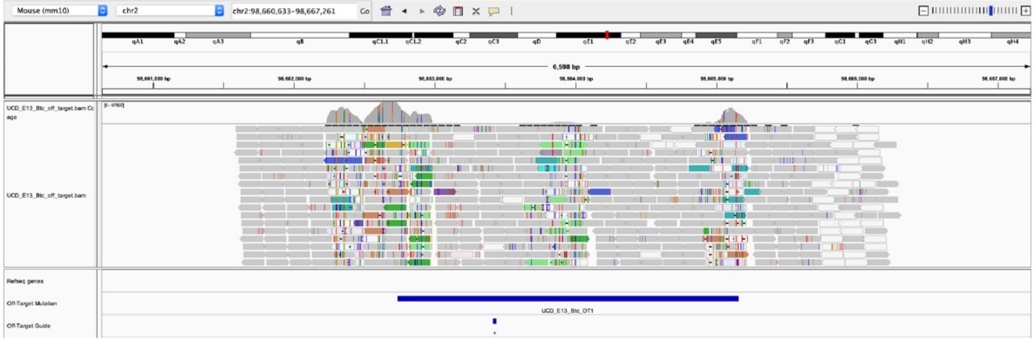

**Supplemental Figure S3**. Whole-genome sequencing evidence for detected Cas9 off-target activity. (**a-z**) IGV screen shots for each region showing coverage, read data, refSeq gene, computationally detected off-target site and position of off-target guide with +/- strand indicated below relative to PAM sequence. Events confirmed by Sanger sequencing contain a green box with ICE results displayed below indicating the detected off-target mutation and estimated percentage of allele contribution. A light blue bar is drawn above the guide sequence and a red bar above the PAM with a vertical dash line indicating Cas9 cut site at the -3 position relative to PAM. Poor sequence quality, low mappability and highly polymorphic regions indicate the that the majority of these are likely false positive results from the software programs and not true off-target events.
